# Expectations and uncertainty shape pain perception during learning

**DOI:** 10.1101/2025.06.24.661242

**Authors:** Mégane Lacombe-Thibault, Michel-Pierre Coll

## Abstract

Pain perception is modulated by expectations and learning processes, but the influence of uncertainty in this relationship is not well established. We aimed to examine the relationship between uncertainty, pain learning and perception using hierarchical Bayesian modeling. In an aversive learning task, fifty participants learned contingencies between auditory cues and painful stimulations under changing levels of uncertainty to create periods of stability and volatility. Model-free analysis of our data suggested unexpected trials resulted in reduced accuracy and greater response times. In unexpected trials, high pain perception was reduced, while low pain perception was increased, in line with documented effects of expectations on pain perception. Computational model fitting revealed participants’ learning was best described by a two-level hierarchical gaussian filter model, suggesting participants adapted their beliefs at multiple levels during the task. Uncertainty influenced pain perception in opposite patterns for high and low pain stimulations: high pain perception was greater under high levels of uncertainty, while there was a non-significant trend for low pain perception to be reduced. Analyses of individual differences suggested depressive symptoms were associated with a reduced learning rate throughout the task. These results shed light on processes involved in pain learning in changing environments. They also suggest a possible relationship between learning alterations and psychological traits commonly found in chronic pain, such as depressive symptoms.

**Perspective:** This article explores the influence of varying levels of uncertainty on pain perception and learning. Findings reveal that uncertainty modulates pain perception differently depending on pain intensity and that pain learning is influenced by psychological traits. These results contribute to our understanding of pain modulation in dynamic environments.

## Introduction

Pain is a fundamental biological signal that serves crucial protective and learning functions^1,2^. While its immediate role in alerting us to potential tissue damage is well understood, pain’s complex relationship with learning and uncertainty remains less clear. In volatile environments, where conditions change unpredictably, learning from pain signals becomes particularly important. However, how uncertainty shapes this process and influences pain perception remains poorly understood^3,4^.

Recent research suggests a fundamental link between pain perception and its learning function. Pain signals appear to carry sensory information and learning signals as prediction errors, which help update future expectations of pain^5–7^. This dual role of pain aligns with both sensory inference and reinforcement learning frameworks^8^. From a sensory inference perspective, pain perception integrates incoming nociceptive signals with prior expectations, allowing for flexible updating of pain predictions^9,10^. Simultaneously, reinforcement learning mechanisms use pain to modify behaviour, with prediction errors serving as teaching signals that guide future actions and expectations^6,11^. The degree to which pain carries new information modulates its perception by increasing or decreasing pain depending on the context^3,12^. While these findings suggest that pain perception is not merely a passive sensory experience but rather an active process combining sensory information with learned expectations, how uncertainty influences this integration remains poorly understood.

Previous studies have examined how uncertainty influences pain perception by experimentally manipulating uncertainty through various approaches. Some varied the probability of receiving pain^13,14^, while others manipulated the predictability of pain intensity or timing^4,15–18^. These manipulations have yielded mixed results: some research suggests that uncertainty increases pain perception^13,14,16^, while others indicate that uncertainty interacts with expectations to influence pain experience^15,17,18^. However, these studies typically employed static manipulations of uncertainty, where participants were explicitly informed about probabilities or experienced fixed levels of unpredictability. Few studies have examined how people learn about changing levels of uncertainty in pain contexts, or how it affects pain perception. An exception is the work of Taylor and collaborators^12^, who found that learned uncertainty was associated with increased pain; however, they employed a task with limited changes in contingencies and thus, limited uncertainty. Consequently, previous research offers limited insight into the relationship between pain and uncertainty, particularly since pain experiences frequently occur in environments where uncertainty fluctuates, necessitating continuous learning^8,10^.

Hierarchical Bayesian models provide a powerful framework for understanding how uncertainty influences pain learning. Unlike traditional associative learning models with fixed learning rates, these models capture multiple forms of uncertainty that evolve dynamically. The Hierarchical Gaussian Filter (HGF), for instance, represents uncertainty from immediate uncertainty about sensory outcomes to higher-order uncertainty about environmental volatility^19,20^. This hierarchical structure allows modeling of different types of uncertainty in pain contexts—from uncertainty about immediate pain intensity to uncertainty about how pain-cue relationships change over time^21,22^. These models have successfully captured learning under uncertainty in various domains and shown, for example, that learned uncertainty is related to stress responses^22^ and illusory pain perception^23^. However, the influence of various forms of uncertainty on pain perception in response to noxious stimuli has been limited.

Using hierarchical Bayesian modeling, we integrated principles from the sensory inference and reinforcement learning frameworks to assess how pain perception is influenced by both sensory prediction errors and uncertainty. We hypothesized that uncertainty would increase pain perception, as pain signals carry greater informational value in volatile contexts requiring continuous updating of predictions. In a task where participants learned associations between auditory cues and painful stimulations under varying levels of uncertainty, we found that estimation uncertainty—reflecting uncertainty about stimulus-outcome relationships—differentially affected pain ratings depending on stimulus intensity. We additionally found that depression was associated with slower updates of pain-cue associations.

## Methods

### Participants

We aimed to recruit 50 participants based on feasibility and to obtain a sample size similar to previous studies using a similar task^24,25^. This sample size provides adequate power (1-β = 0.80) to detect a moderate relationship of |r| = 0.37 with a significance threshold of α = 0.05, two-tailed, for our model-free analysis. For Bayesian model comparison, the sample size of 50 participants each completing 192 trials of the learning task was considered sufficient based on previous studies using similar tasks. Fifty-two participants with no history of neurological, psychiatric or pain disorders took part in the experiment. Two participants were excluded for misidentifying a high number of stimulations (accuracies of 73.9% and 77.6%, z scores of -4.11 and -3.44 relative to the group average of 97%). The analyses were performed on the remaining 50 participants which included 28 participants who self-identified as cisgender women and 22 as cisgender men (mean age=24.38 years; SD=4.27). Forty-three participants identified as White (86%), 2 as Asian (4%), 2 as Latino/a American (4%), 1 as Black (2%), 1 as Middle Eastern (2%) and 1 as North African (2%). To encourage diversity and inclusion in our sample, we employed a broad recruitment strategy that included emails sent to the entire university community as well as posters in various locations across campus and within the research center. No members of the public or patients were involved in this study’s design or implementation.

Written informed consent was obtained for all participants, and this experiment was approved by the ethics committee of the *Centre intégré universitaire de santé et de services sociaux de la Capitale-Nationale* (project #2022-2452) and participants received $20 Canadian dollars for their participation.

### Electrical stimulation

Painful electrical stimuli were administered using a biphasic constant current stimulator (DS8R, Digitimer Ltd, United Kingdom) combined with a train delay generator (DG2A, Digitimer Ltd, United Kingdom) to deliver, at each trial, 5 monophasic stimulations of 2 ms each presented at 200 Hz. Electrical stimuli were delivered through two 8mm Ag/AgCl cup electrodes filled with conductive gel and placed 2 centimetres apart on the back of the participants’ non-dominant hand. Prior to electrode placement, the hand was cleaned with alcohol swabs.

### Procedure

The experimental session lasted approximately 80 minutes and was divided into 3 phases: 1) Calibration, 2) Aversive learning task and 3) Self-reported questionnaires.

### Calibration

Pain levels for the experiment were determined using a staircase method^26^. The stimulation intensity was increased from 1 mA in 1 mA increments until participants indicated they had reached their pain tolerance threshold. Participants were informed a stimulation similar to their pain tolerance threshold could be used during the task, where they would receive it repeatedly. They were asked to keep this in mind when selecting their maximal stimulation during the calibration process. After each stimulation, participants rated the intensity of the painful stimulation on a visual analog scale (VAS). The VAS was labelled from “no pain” to “worst imaginable pain”. Participants were also asked after each stimulation if they were willing to receive a more intense stimulation.

This ramping procedure was repeated 3 times to ensure participants were familiar with the electrical stimulations and rating scale. Using the results from the third ramp, the intensity at which participants first reported perceiving pain (VAS rating > 0) was defined as the pain detection threshold, while the maximum tolerable intensity was defined as the pain tolerance threshold, which was not exceeded during the rest of the experiment.

The second phase of the calibration consisted of 14 equidistant electrical stimulations between the pain detection and tolerance thresholds presented in a random order. Participants rated the intensity of each stimulation using the same VAS as in the first part of the calibration. This process aimed to avoid expectation effects on pain perception.

The intensities corresponding to 40% (low pain) and 90% (high pain) of the maximum pain rating were determined by fitting various polynomial functions to the relationship between each stimulation intensity and the corresponding VAS rating obtained during the second phase of the calibration. The function that best represented this relationship was used to identify the low and high painful stimulations used for the rest of the experiment. On average, the intensity of the low pain stimulation was 4.78 mA (SD=2.57, range: 1.5 mA to 13 mA) and the intensity of the high pain stimulation was 8.93 mA (SD=4.02, range: 4 mA to 21 mA) with an average difference of 4.21 mA (SD=1.89, range:1.25 mA to 10.5 mA) between the two stimulations. For one participant, high and low pain stimulation intensities were reduced after the first block of the task, as the calibrated intensities were not well tolerated. Reduced intensities are reported in the descriptive statistics above.

### Aversive learning task

The experimental task was based on a previously used learning paradigm in the context of volatility^25,27^ and lasted approximately 20 minutes. It included 192 trials organized into 4 blocks with a brief pause following each block. The task began with 4 practice trials so participants could familiarize themselves with the task.

In each trial (Figure 1a), a fixation cross appeared on the screen for a random duration between 1 and 2 seconds, followed by a low (330 Hz) or a high (660 Hz) tone presented for 300 ms. The sounds were presented at a clearly audible intensity and were easily distinguishable by participants, as verbally confirmed by participants prior to the task. Two hundred ms after the end of the tone, a low or high intensity painful electrical stimulus was delivered for 30 ms. With their dominant hand, participants then had to indicate as quickly as possible within the two-second time limit whether they received the low or high intensity stimulation by pressing one of two keys on a response box. At the end of each trial, participants had up to 3 seconds to rate the intensity of the painful stimulus on the same VAS as used in the calibration. VAS position was converted to a 0 to 100 score for analyses. If participants exceeded the allowed time at each step of the task, the trial ended and was not considered in analyses.

**Figure 1.**
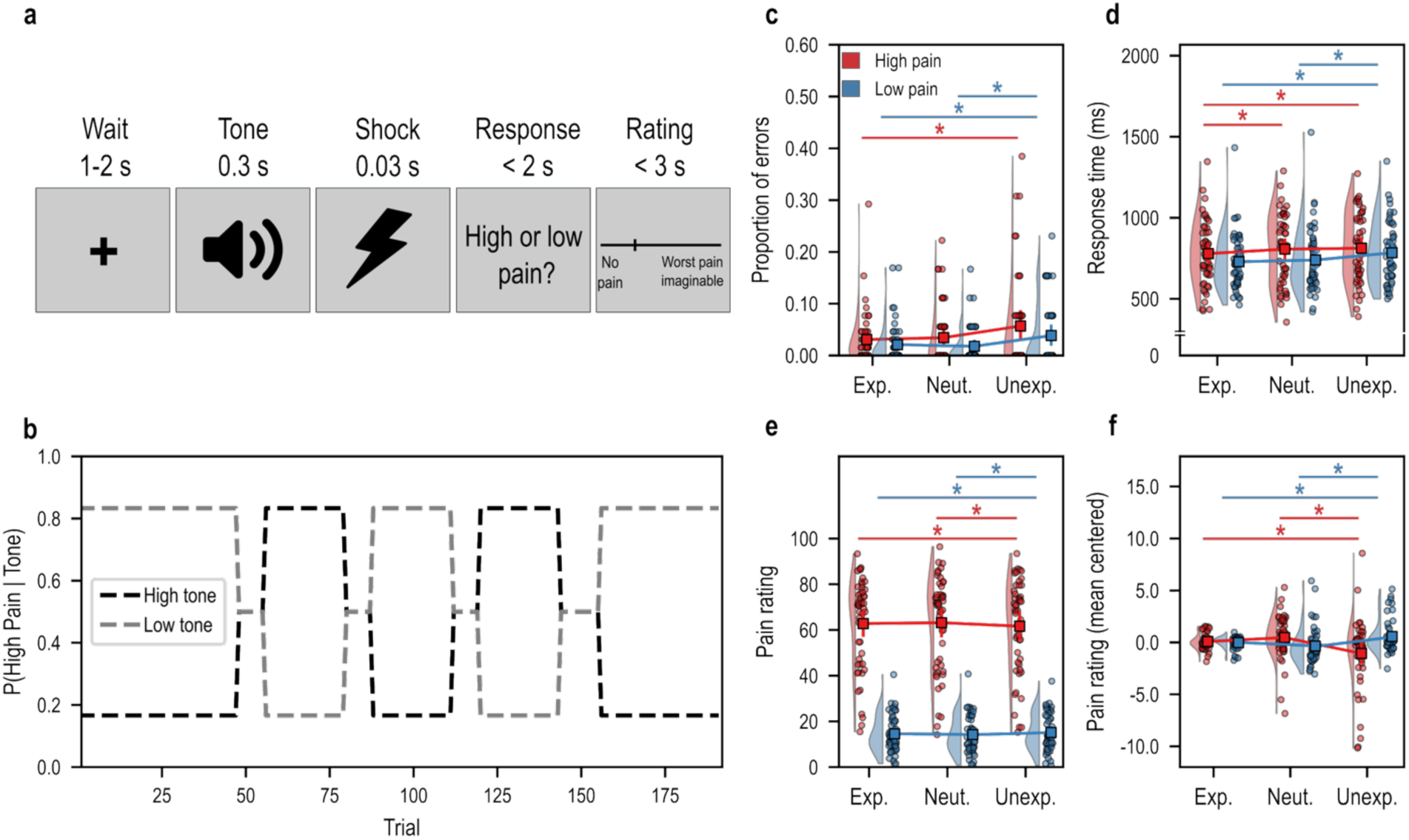
(a) Schematic representation of an experimental trial. (b) Tone and pain level contingencies throughout the experimental task showing phases in which the high or low pain level could be expected (Exp., 0.84), Neutral (Neut. 0.5) or Unexpected (Unexp., 0.16). (c) Proportion of errors in identifying the pain levels, (d) response time (e) pain rating and (f) pain rating centered on each pain level mean as a function of pain level and contingencies. All error bars show the 95% confidence interval of the mean.

Although participants were not informed of the exact task structure and pattern of tone-pain association changes shown in Figure 1b, they were aware that the tone frequency was associated with the probability of receiving low or high intensity painful stimulations. The tone-pain probability structure was predetermined and changed every 8, 12, 24, 36 or 48 trials (blocks), creating periods of stability and volatility (Figure 1b). Within each block, trials were pseudo-randomly presented to match the block’s length and probability structure. During each block, the probability of receiving high or low pain after a given tone could be strong (probability=0.84), while the pain-intensity probability for the other tone was weak (probability=0.16), or both probabilities could be null (probability=0.5). For model free analyses, we used the same labels as previous studies^25,27^ and we label these outcomes as expected (P(pain level|tone) = 0.84), unexpected (P(pain level|tone) = 0.16) and neutral (P(pain level|tone) = 0.5) to reflect the outcome probabilities defined by the task structure.

The order of stimuli presentation was adjusted so that in each block, both tones appeared as frequently and were presented in a random order. Throughout the task, the probability of one outcome (e.g., low pain) given a cue (e.g., low tone) was the same as the probability of the other outcome (e.g., high pain) given the other cue (e.g., high tone). These manipulations ensured that the probability of receiving low or high pain before the presentation of the sound was 50% on any given trial^24,27^. This ensured a similar distribution of high and low pain stimuli across blocks.

### Self-reported questionnaires

After the experiment, participants completed the French versions of the Pain Catastrophizing Scale (PCS^28,29^), the State-Trait Anxiety Inventory (STAI^30,31^), and the Beck Depression Inventory-II (BDI-II^32^). These questionnaire data were used in exploratory analyses to examine how psychological traits might influence aversive learning under uncertainty.

### Computational modeling

To model participants’ learning throughout the task, we compared several computational models: a Rescorla-Wagner (RW) model, a Pearce-Hall (PH) model and the Hierarchical Gaussian Filter (HGF). For the HGF, we tested four variants that differed in their complexity: models with either two or three levels, each implemented with or without a perceptual uncertainty parameter.

For the learning models, pain outcomes at each trial were coded in contingency space to account for the tone-pain associations, as recommended in the HGF toolbox^19,20^ and previous studies using a similar task^25,33^. Specifically, high or low pain following low tones were coded 1 and high or low pain following high tones were coded 0, therefore reflecting the contingency between cue (tones) and outcome (pain intensity). These binary outcomes served as observed sensory inputs in the models, capturing both the cue presented and the resulting sensory outcome. Using these outcomes as observed sensory inputs, we assessed each model’s ability to predict participants’ log response times using a linear response model. We used the default implementation of each learning model in the TAPAS toolbox^34^ and we thus refer readers to the open code (https://github.com/translationalneuromodeling/tapas, version 7.1.2) for the update equations of each model. The required priors for the HGF models were adjusted from the default values defined in the TAPAS toolbox based on the priors used in a previous study using a similar task^25^ (see Supplementary Table S1 for all priors for the HGF models). Trials with mistakes or response times below 200 ms were ignored in the response model fitting procedure (average number of trials included = 186.22, SD=7.38, range = 154-192).

The models were fitted to the data using the HGF module of the TAPAS toolbox^34^. Model fits at the group level were compared within a Bayesian model comparison procedure, as implemented in the Variational Bayesian Analysis toolbox (VBA; https://mbb-team.github.io/VBA-toolbox). Specifically, model fit was quantified using the log model evidence (LME) for each participant, which represents the negative surprise of the data under a given model^19^ and offers a trade-off between model fit and complexity^25^. Model selection was done by performing group-level estimations of the frequency of a given model being the best across participants and the protected exceedance probability of each model, representing the probability that a given model is more likely than others^35^. The model with the highest exceedance probability and most frequently identified as the best model was considered the winning model.

Below, we briefly describe each model and the response models linking the trial-wise estimates derived from these models to the log of the response time at each trial.

### Rescorla-Wagner model

The RW model uses prediction errors to drive associative learning: larger discrepancies between actual and expected outcomes produce greater changes in associative strength^36^. At each trial (t), the associative strength (V) between auditory cues and pain outcomes is updated based on a learning rate parameter (ɑ) and the prediction error (δ)—the difference between the actual outcome (λ) and expected outcomes. The model assumes that response times are predicted by the trial-specific absolute prediction error (|δ|), with five free parameters: the learning rate (ɑ), initial associative strength (V0) and the response model coefficients and noise (β₀, β₁, ζ).

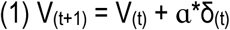

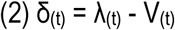

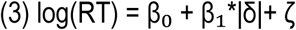

### Pearce-Hall model

The Pearce-Hall model has previously been used in uncertainty studies^12,37^ and combines associative learning with dynamic attention. The model assumes that response times were influenced by attention (α) at each trial (t). The attention parameter is updated based on the absolute prediction error from the previous trial, while associative strength (V) follows the Rescorla-Wagner learning rule. This model included five free parameters: initial attention (α₀), attention learning rate (η), three response coefficients (β₀, β₁), plus Gaussian noise (ζ), as specified in the linear response model below:

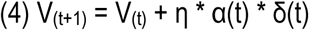

Where

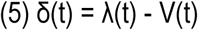

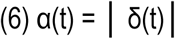

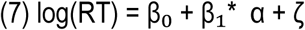

### Hierarchical gaussian filter

The Hierarchical Gaussian Filter model (HGF; Figure 2a-b) assumes that response times were influenced by uncertainty estimates across two (HGF2) or three (HGF3) hierarchical levels. The model tracks belief updates about stimulus transitions at two or three coupled levels: stimulus outcomes (x₁), probability of stimulus transitions (x₂), and, for the three-level version, volatility of these probabilities (x₃). The beliefs (µ) at each level above the first evolve as Gaussian random walks, where ω₂ determines the step size of probability updates at the second level, and in the HGF3, ω₃ determines the step size of volatility updates at the third level. Each level is updated based on precision-weighted (𝜓) prediction errors (𝜀, 𝛿) from the level below. The equations can be found in previous descriptions of the model^19,20^.

**Figure 2.**
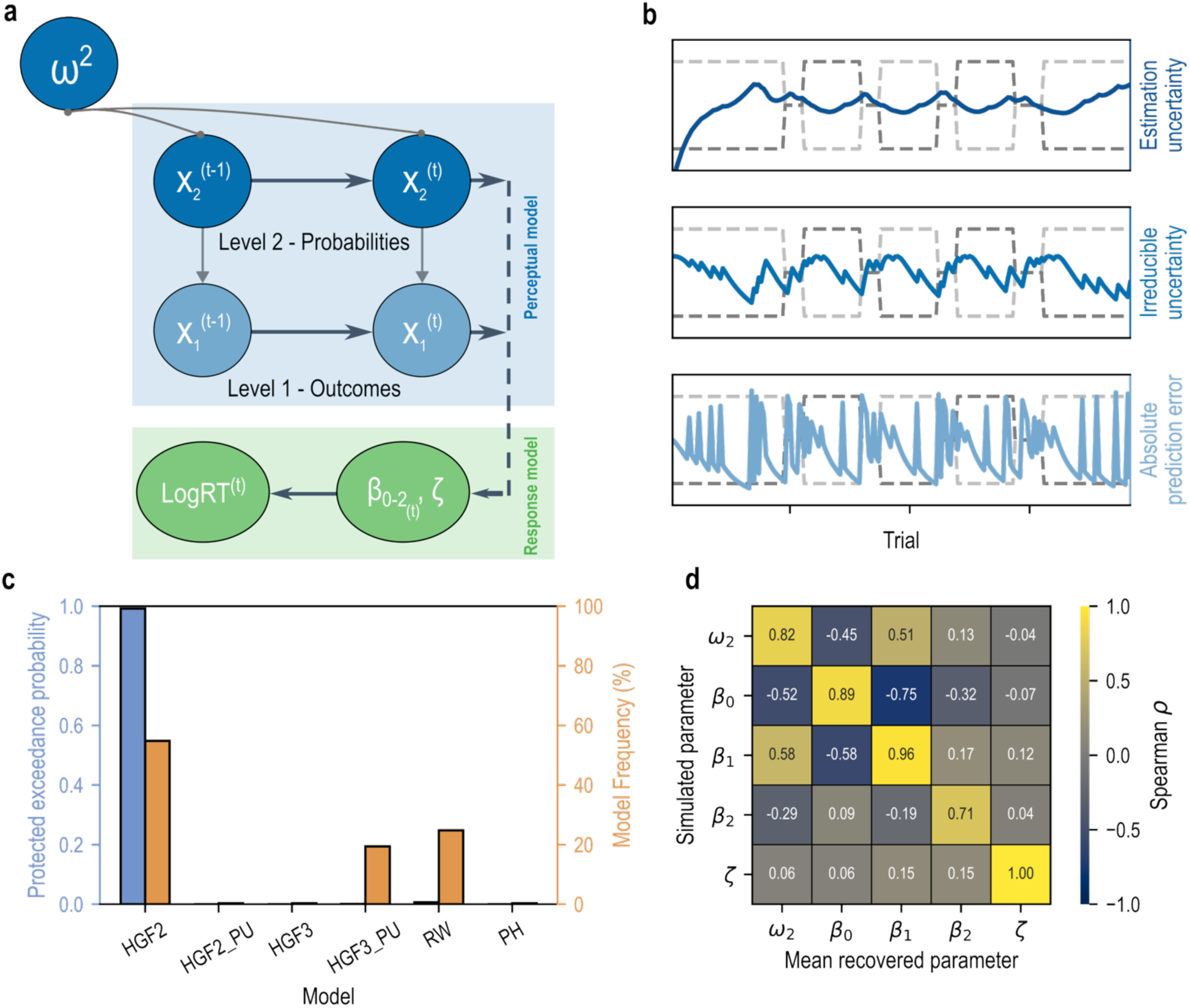
(a) Structure of the HGF2 model. (b) Example of absolute prediction error, irreducible and estimation uncertainty across trials. (c) Results of the Bayesian model comparison. (d) HGF2 model parameter recovery.

For both HGF2 and HGF3, we tested versions that took into account perceptual uncertainty in pain processing (HGF2pu and HGF3pu). In these versions, rather than treating outcomes as binary events, the model represents them as continuous values drawn from a mixture of Gaussian distributions. The perceptual uncertainty parameter α represents the variance of these distributions centered at 0 and 1, with higher values indicating greater difficulty in discriminating between high and low pain intensities^19^.

The HGF linear response model incorporated uncertainty estimates from each hierarchical level transformed to the outcome level^24,25^: outcome uncertainty (unc₁), estimation uncertainty (unc₂), and, for the HGF3, phasic volatility (unc₃). The number of free parameters varied across HGF variants: five to eight parameters depending on model complexity. For the learning component, these included step size parameters (2^nd^ level tonic volatility; ω₂, and for HGF3, 3^rd^ level volatility ω₃) and, where applicable, perceptual uncertainty (α). The response model included coefficients for each uncertainty term (β0-2 for HGF2, β0-3 for HGF3) and Gaussian noise (ζ):

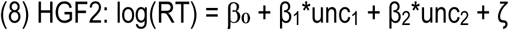

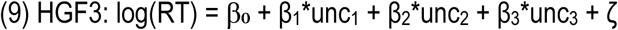

### Statistical analyses

#### Model-free analyses

We first confirmed the influence of the variations in contingencies throughout the task on behavioural responses independently of the computational modeling procedure. To this end, we used repeated-measures analyses of variance (ANOVA) to test the effect of Trial type (expected [84% probability], unexpected [16% probability] or neutral [50% probability]) and Pain intensity (high or low) and their interaction on response times (RTs), proportion of errors and pain intensity ratings. The Greenhouse-Geisser correction was used in case of violation of the sphericity assumption and post-hoc pairwise comparisons were conducted. The level of significance was set at *p*<0.05, bilateral, for all analyses.

#### Model-based analyses

We used linear mixed-effects models to examine how trial-by-trial pain ratings were influenced by computational estimates from the winning model, the HGF2: the absolute prediction error at the first level (\delta_1), outcome uncertainty (\hat{\sigma_1}) and estimation uncertainty (\hat{\sigma_2}). For each computational estimate, we constructed a separate model that included the estimate, stimulation intensity (high vs. low), and their interaction as fixed effects. All models included random intercepts and random slopes for participants for all fixed effects, accounting for individual differences in baseline ratings and in the strength of relationships between predictors and pain ratings. Estimation uncertainty and outcome uncertainty were respectively log transformed and squared to take into account their positive and negatively skewed distributions and favour model convergence.

#### Individual differences in learning, pain catastrophizing, depressive and anxiety symptoms

The individual differences in the PCS, BDI and STAI-Y1 and STAI-Y2 questionnaires total scores were examined by correlating these values with the parameters of the winning learning model. Pearson r correlations were conducted with the data from 49 participants only, as one participant had missing data on the questionnaires due to a technical issue. The Holm-Bonferroni method was used to correct alpha tests.

## Results

### Model-free analyses

We confirmed that Trial type influenced both participants’ accuracy [F(1.644,80.542)=7.423, p=0.002, η_p_^2^=0.030; Figure 1c] and response times [F(1.908,93.508)=8.272, p<0.001, η_p_^2^=0.008; Figure 1d]. As expected, participants made more errors and were slower in unexpected trials compared to both expected [accuracy: t(49)=3.101, p=0.003, Hedge’s g=0.414; RT: t(49)=-3.955, p<0.001, g=-0.236] and neutral trials [accuracy: t(49)=2.921, p=0.005, g=0.420; RT: (t(49)=-2.137, p=0.038, g=-0.128]. Participants also made more errors [(F(1,49)=5.027, p=0.030, η_p_^2^=0.016, *g*=-0.324)] and were slower [F(1,49)=12.287, p<0.001, η_p_^2^=0.015)] in high pain trials compared to low pain trials. There was no interaction between Trial type and Pain intensity on accuracy [F(1.627,79.697)=0.248, p=0.735, η_p_^2^<0.001] or RTs [F(1.677,82.169)=2.474, *p*=0.100, η_p_^2^=0.002].

We further confirmed that pain intensity ratings differed between the two administered intensities (F(1,49)=315.539, p<0.001, η_p_^2^=0.707; Figure 1e). Pain intensity and Trial type showed an interaction (F(1.575,77.176)=8.259, p=0.001, η_p_^2^=0.001): participants rated high pain stimulations lower during unexpected trials compared to both expected (t(49)=2.084, p=0.042, g=0.056) and neutral trials (t(49)=2.400, p=0.020, g=0.073), while ratings for low pain trials increased in unexpected conditions compared to both expected (t(49)=-2.151, p=0.036, g=-0.071) and neutral trials (t(49)=-2.616, p=0.012, g=-0.111; Figure 1f).

### Computational model fitting and validation

Bayesian model comparison identified the two-level HGF without perceptual uncertainty (HGF2) as the best predictor of participants’ log response times (Figure 2c; protected exceedance probability [PEP] = 0.99; model frequency = 0.540).

To validate this model, we conducted two analyses. First, we performed parameter recovery by simulating ten response time datasets for each participant using their fitted HGF2 parameters (500 datasets total) and generating response times by adding random gaussian noise to the linear response model. We then fitted the model to these simulated data and compared the recovered parameters, averaged across the ten datasets per participant, to the original simulation parameters. This analysis revealed high Spearman ρ correlations between simulated and recovered parameters (Figure 2d; parameter relationships illustrated in Supplementary Figure S1c-g).

Second, we assessed model recovery by fitting all models to data simulated from each model. The HGF2 was correctly identified as the winning model for 95% of datasets simulated using its parameters with no evidence for better fit from alternative models. However, the HGF2 was not specific in recovering itself and showed high recovery for datasets generated using other models (Supplementary Figure S1b), further suggesting that the HGF2 was a superior fit to our task.

### Model-based analyses

Analysis of the relationship between the HGF2 model’s trial-wise estimates and pain ratings revealed a significant interaction between estimation uncertainty and pain intensity (β = 8.194, SE = 2.191, p < 0.001; Figure 3a). Decomposition of this interaction revealed that estimation uncertainty was positively associated with pain ratings in high-intensity trials (β = 5.669, SE = 1.891, p = 0.003), while showing a negative but not significant relationship with pain ratings in low-intensity trials (β = -2.702, SE = 1.385, p = 0.05). Similarly, there was a significant interaction between absolute prediction errors and pain ratings (β = -2.994, SE = 1.437, p = 0.037; Figure 3b). The decomposition of this interaction revealed that absolute prediction errors were negatively associated with pain ratings in high pain trials (B = -2.264, SE = 1.122, p = 0.044) but showed no significant relationship in low pain trials (B = 0.731, SE = 0.664, p = 0.271). Outcome uncertainty did not significantly affect pain ratings (β = -3.540, SE = 10.952, p = 0.747), and its interaction with pain intensity was also non-significant (β = -1.221, SE = 24.563, p = 0.960).

**Figure 3.**
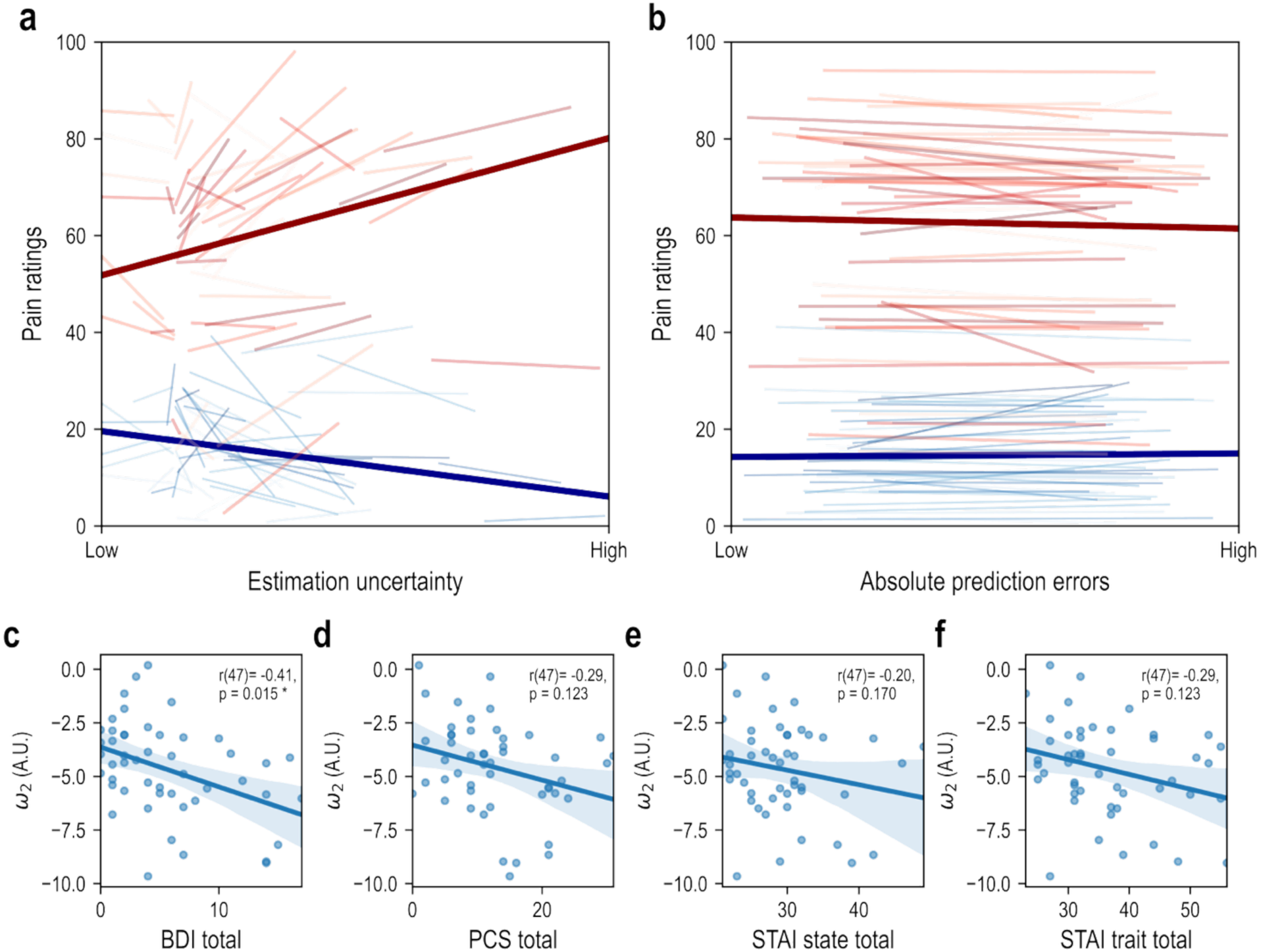
Model-based analyses results of linear mixed-effects models. (a) Pain ratings by estimation uncertainty where lighter lines represent each participant’s individual slope and darker bold lines represent the average slope (red for high pain intensity and blue for low pain intensity). (b) Pain ratings by absolute prediction errors. (c-f) Individual differences in BDI, PCS, STAI state and STAI trait scores and the tonic volatility parameter (ω₂). The reported p-values were corrected with the Holm-Bonferroni method.

### Individual differences in learning

High depressive symptoms were associated with lower tonic volatility for tone-pain associations (ω₂; r(47) = -0.41, p = 0.015; Figure 3c). Pain catastrophizing scores (r(47) = -0.29, p = 0.123; Figure 3d), state anxiety (r(47) = -0.20, p = 0.170; Figure 3e) and trait anxiety (r(47) = -0.29, p = 0.123; Figure 3f) showed non-significant negative correlations with tonic volatility.

## Discussion

Pain perception is fundamentally shaped by expectations and learning. Using a dynamic learning task, we asked participants to learn associations between auditory cues and painful electrical stimulations of two different intensities. The task was designed to generate periods of high and low uncertainty by frequently changing the cue-pain contingencies throughout the experiment. Results of computational modeling revealed that under high uncertainty, high-intensity pain was perceived as more painful, while low-intensity pain was perceived as less painful. These findings extend previous research on expectation and uncertainty effects in pain perception^4,12,18,38^ by showing how learned uncertainty in a changing environment modulates pain processing in an intensity-dependent manner—amplifying the perception of high pain while attenuating the perception of low pain.

Model-free analyses showed expected results, as more errors and longer response times were found in unexpected trials. Our results also suggest an opposite effect of expectations on pain perception for high and low intensity stimuli, where high pain perception decreased and low pain perception increased in unexpected trials. These findings are in line with Bayesian inference theories where perception is biased toward expectations^39^, as found in similar studies^18^. Although these analyses provide insight on our findings, they fail to explain the underlying mechanisms responsible for the effect of unexpectedness. Computational modeling allows for a more in-depth explanation of the dynamic relationships between expectations and uncertainty and how they influence perception to optimize learning.

Our computational modeling approach revealed that learning about pain in volatile environments is best captured by hierarchical learning processes. The two-level Hierarchical Gaussian Filter (HGF2^19,20^) outperformed other models in predicting participants’ response times, suggesting that participants continuously updated beliefs at multiple levels. Similarly to a recent study^23^, the three-level HGF3, which additionally models estimates of environmental uncertainty, did not outperform the simpler two-level HGF and showed poor parameter recovery performance in comparison to the HGF2. This discrepancy, when compared to several other studies using a similar task^24,25^, could be attributed to the specific contingency structure of our task, which included fewer trials and contingency switches as well as a repetitive contingency switching pattern compared to a random one, therefore not requiring estimations of volatility to perform adequately.

Using the HGF2 to estimate participants’ subjective perception of environmental uncertainty, we found that its relationship with pain perception depended on stimulus intensity: under higher estimation uncertainty, high-intensity pain was perceived as more intense, while there was a trending negative relationship for low-intensity pain, suggesting it was perceived as less intense, though this effect did not reach statistical significance. This pattern aligns with Bayesian frameworks of perception and suggests that under uncertainty, the pain system shifts toward relying more on incoming sensory information^39^. It could also be the case that the differential impact of uncertainty on pain perception depending on stimulus intensity reflects asymmetric cost functions for pain perception and serves to optimize pain’s learning and control function^6^. When facing high pain stimuli under uncertainty, the biological cost of underestimating the threat outweighs the cost of overestimating, leading to elevated pain ratings as a protective learning mechanism for avoiding significant threats. Conversely, for low pain stimuli, the system favors underestimation when uncertain, as this minimizes costs associated with unnecessary defensive responses, resulting in lower pain ratings. This could optimize learning by filtering out noisier signals of unthreatening pain. Our results could also explain the previously observed mixed effects of uncertainty on pain perception^13,14,16–18^, as studies using different pain intensities might find apparently contradictory effects of uncertainty depending on where their stimuli fell relative to participants’ average expectations. Here, our dynamic modulation of uncertainty through changing contingencies provides a more ecologically valid approach compared to previous studies that used static manipulations of uncertainty or fixed probabilities.

Additionally, we found absolute prediction errors, as estimated by the HGF2, influenced pain perception differently according to stimulus intensity: high pain was rated lower as prediction errors increased, while there was evidence for low pain to be rated higher, although this effect did not reach statistical significance. These results align with learning theories of pain suggesting that perception is dynamically adjusted in response to prediction errors to optimize learning^6^. The greater malleability of high pain compared to low pain could reflect an asymmetric cost of enhancing the salience of prediction errors. Less threatening signals of pain could carry less informational value for harm avoidance, thus requiring less adjustment in response to prediction errors. In contrast, more salient signals of high pain could be more flexible in response to prediction errors and serve a greater learning purpose for threat avoidance.

Individual differences in self-reported measures of depression, anxiety and pain catastrophizing were examined in relation to learning parameters estimated by the HGF2 model. We found a non-significant negative trend between anxiety and pain catastrophizing and ω₂—a parameter representing the step size of the learning rate for updating cue-outcome associations over time^19^—while participants reporting more symptoms of depression showed significantly reduced ω₂ values. A lower ω₂ indicates reduced flexibility in updating beliefs about how pain-cue relationships change. This finding has particular relevance for chronic pain populations, where elevated catastrophizing, depression and anxiety are common^40^ and represent important risk factors for chronic pain development^41^. Our results suggest that these traits may be linked with impaired ability to update pain expectations in changing environments, potentially contributing to maladaptive pain processing. It should be noted that the three measures were moderately to highly correlated (r = 0.5 to 0.72), suggesting that they tap into a common trait related to learning. Nevertheless, the relationships between these traits and learning may help explain why some individuals develop chronic pain conditions while others recover from similar injuries and thus provide a computational hypothesis for the long-theorized link between chronic pain and learning mechanisms^42,43^ that should be tested in future studies.

Several limitations warrant consideration. First, while we used a similar design to previous studies^44,45^, our task employed a single contingency structure. Our findings should be tested for generalization across different types of contingency patterns and learning environments. Second, while our findings suggest a link between psychological characteristics and learning mechanisms, these relationships were relatively weak and observed in a healthy sample with few individuals reporting high anxiety, depression or catastrophizing.

Testing these relationships in clinical populations will be crucial to establish their relevance to chronic pain. Finally, while we apply our results to pain perception, similar influences of uncertainty on perception might be observed in other phenomena, and psychological traits might be linked to general learning mechanisms rather than being specific to pain learning.

In conclusion, our study demonstrates that pain perception is dynamically modulated by learned uncertainty, with distinct effects depending on pain intensity, and is influenced by individual differences in psychological characteristics. The successful application of hierarchical Bayesian models to capture these effects provides a promising framework for understanding pain learning in complex, changing environments. Our finding that symptoms of depression are associated with reduced learning efficiency could be particularly relevant for understanding how individuals with chronic pain learn and deal with uncertainty in their daily lives.

## Disclosures

MLT was supported by scholarships from the *Fonds de Recherche Québec – Santé* (2023-2024, 2024-2028) and the Canadian Institutes of Health Research (2023, 2025-2028). MPC and this work was supported by a career grant from the *Fonds de Recherche Québec - Santé* (grant #309041, 2021-2025). The authors declare that they have no conflict of interest. All raw data and code necessary to reproduce all analyses and figures are available at https://github.com/meganel-t/painuncertainty.

## Notes

https://github.com/meganel-t/painuncertainty

### Competing Interest Statement

The authors have declared no competing interest.

### Summary of Updates

Introduction and Methods sections have been updated to clarify the experimental design and coding scheme. Figures have been revised.

