## Supplementary Materials for "Expectations and uncertainty shape pain perception during learning"

**Table S1. Priors for the different HGF perceptual and response models used during model fitting.**

| **Parameter name** | **Prior mean** | **Prior variance** |
| --- | --- | --- |
| **Perceptual model** |  |  |
| Tonic volatility (level 2: ω_2_) | -5 | 16 |
| Tonic volatility (level 3: ω_3_) | -6 | 16 |
| Perceptual uncertainty (α) | log(0.5) | 1 |
| **Response model** |  |  |
| Baseline RT (𝛽_0_) | log(760) | 4 |
| RT predictors (𝛽) | 0 | 4 |
| Decision noise (𝜁) | log(log(20)) | log(2) |

**S1.2. Model recovery**

Model recovery performance for other models was also compared and suggests the RW model (proportion = 44%) had the best recovery performance while the HGF3pu model (proportion = 28%) had moderate recovery performance (Figure S1). All other models showed poor model recovery performance. Analyses of model confusion suggest the HGF2 model was consistently mistaken with the HGF2pu (proportion = 100%), HGF3 (proportion = 78%), HGF3pu (proportion = 64%), RW (proportion = 56%) and PH models (proportion = 90%; Figure S1a). Protected exceedance probability (PEP) analyses suggested the HGF2 model clearly outperformed itself as well as the HGF2pu, HGF3, HGF3pu, RW and PH models (PEPs = 1.00; Figure S1b).

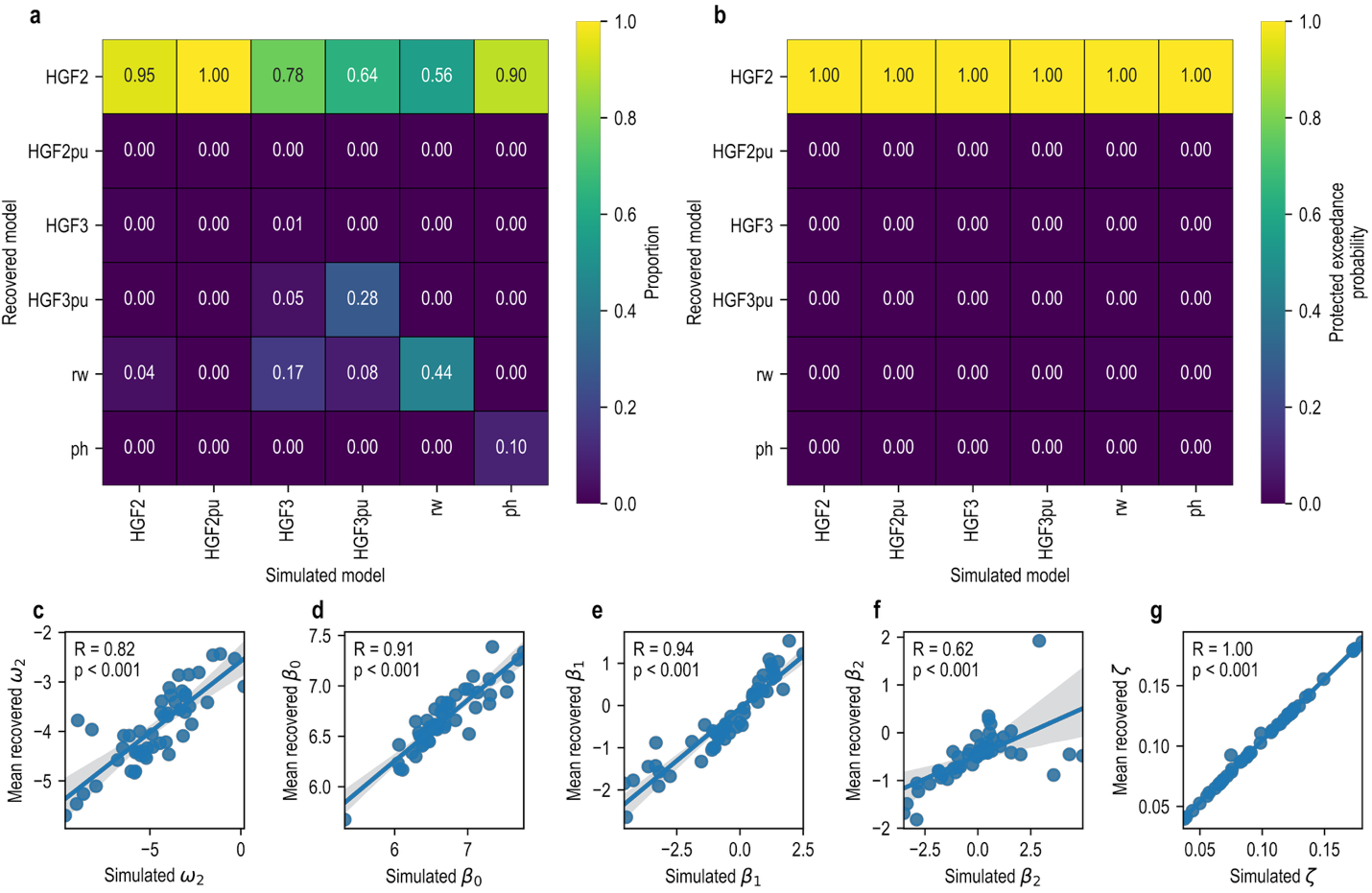

**Figure S1.** (a) Proportion and (b) protected exceedance probability of simulated and recovered models. (c-g) Spearman ρ correlations between simulated and mean recovered parameters of winning models (HGF-3pu).

**S1.3. Full results for statistical analyses of model-free results**

| **Proportion of errors** | |
| --- | --- |
| **3 (trial type) x 2 (stimulus intensity) ANOVA** | |
|  | **Statistics** |
| **Trial type** | F(1.644,80.542)=7.423, *p*=0.002, η_p_^2^=0.030 |
| **Stimulus intensity** | F(1,49)=5.027, *p*=0.030, η_p_^2^=0.016 |
| **Trial type x stimulus intensity** | F(1.627,79.697)=0.248, *p*=0.735, η_p_^2^=0.001 |
| **Pain ratings** | |
| **3 (trial type) x 2 (stimulus intensity) ANOVA** | |
|  | **Statistics** |
| **Trial type** | F(1.696,83.078)=0.632, *p*=0.509, η_p_^2^<0.001 |
| **Stimulus intensity** | F(1,49)=315.539, *p*<0.001, η_p_^2^=0.707 |
| **Trial type x stimulus intensity** | F(1.575,77.176)=8.259, *p*=0.001, η_p_^2^=0.001 |
| **Response times** | |
| **3 (trial type) x 2 (stimulus intensity) ANOVA** | |
|  | **Statistics** |
| **Trial type** | F(1.908,93.508)=8.272, *p*<0.001, η_p_^2^=0.008 |
| **Stimulus intensity** | F(1,49)=12.287, *p*<0.001, η_p_^2^=0.015 |
| **Trial type x stimulus intensity** | F(1.677,82.169)=2.474, *p*=0.100, η_p_^2^=0.002 |

| **Proportion of errors** | | | | | | |
| --- | --- | --- | --- | --- | --- | --- |
| **Comparison** | | **T** | **df** | **p** | **BF10** | **Hedge’s g** |
| Trial type | Expected v. Neutral | 0.055 | 49 | 0.956 | 0.154 | 0.007 |
|  | Expected v. Unexpected | 3.101 | 49 | 0.003 | 10.152 | 0.414 |
|  | Neutral v. Unexpected | 2.921 | 49 | 0.005 | 6.557 | 0.420 |
| Stimulus intensity | Low v. High | -2.242 | 49 | 0.030 | 1.505 | -0.324 |
| Stimulus intensity (high) v. trial type | Expected v. Neutral | 0.469 | 49 | 0.641 | 0.171 | 0.069 |
|  | Expected v. Unexpected | 2.609 | 49 | 0.012 | 3.209 | 0.347 |
|  | Neutral v. Unexpected | 1.846 | 49 | 0.071 | 0.739 | 0.291 |
| Stimulus intensity (low) v. trial type | Expected v. Neutral | -0.473 | 49 | 0.638 | 0.171 | -0.083 |
|  | Expected v. Unexpected | 2.164 | 49 | 0.035 | 1.296 | 0.333 |
|  | Neutral v. Unexpected | 2.259 | 49 | 0.028 | 1.554 | 0.401 |
| **Pain ratings** | | | | | | |
| **Comparison** | | **T** | **df** | **p** | **BF10** | **Hedge’s g** |
| Trial type | Expected v. Neutral | -0.106 | 49 | 0.916 | 0.155 | -0.002 |
|  | Expected v. Unexpected | 0.960 | 49 | 0.342 | 0.238 | 0.023 |
|  | Neutral v. Unexpected | 0.858 | 49 | 0.395 | 0.218 | 0.025 |
| Stimulus intensity | Low v. High | 17.763 | 49 | <0.001 | 7.379e19 | 3.072 |
| Stimulus intensity (high) v. trial type | Expected v. Neutral | -1.100 | 49 | 0.277 | 0.272 | -0.019 |
|  | Expected v. Unexpected | 2.084 | 49 | 0.042 | 1.118 | 0.056 |
|  | Neutral v. Unexpected | 2.400 | 49 | 0.020 | 2.061 | 0.073 |
| Stimulus intensity (low) v. trial type | Expected v. Neutral | 1.057 | 49 | 0.296 | 0.260 | 0.041 |
|  | Expected v. Unexpected | -2.151 | 49 | 0.036 | 1.265 | -0.071 |
|  | Neutral v. Unexpected | -2.616 | 49 | 0.012 | 3.262 | -0.111 |
| **Response times** | | | | | | |
| **Comparison** | | **T** | **df** | **p** | **BF10** | **Hedge’s g** |
| Trial type | Expected v. Neutral | -1.975 | 49 | 0.054 | 0.919 | -0.101 |
|  | Expected v. Unexpected | -3.955 | 49 | <0.001 | 100.812 | -0.236 |
|  | Neutral v. Unexpected | -2.137 | 49 | 0.038 | 1.232 | -0.128 |
| Stimulus intensity | Low v. High | 3.505 | 49 | <0.001 | 28.855 | 0.253 |
| Stimulus intensity (high) v. trial type | Expected v. Neutral | -2.094 | 49 | 0.041 | 1.138 | -0.131 |
|  | Expected v. Unexpected | -2.266 | 49 | 0.028 | 1.577 | -0.161 |
|  | Neutral v. Unexpected | -0.367 | 49 | 0.716 | 0.164 | -0.027 |
| Stimulus intensity (low) v. trial type | Expected v. Neutral | -0.999 | 49 | 0.323 | 0.246 | -0.057 |
|  | Expected v. Unexpected | -4.203 | 49 | <0.001 | 208.023 | -0.299 |
|  | Neutral v. Unexpected | -2.886 | 49 | 0.006 | 6.029 | -0.227 |
